# Subtypes of functional connectivity associate robustly with ASD diagnosis

**DOI:** 10.1101/2020.04.14.040576

**Authors:** Sebastian G. Urchs, Angela Tam, Pierre Orban, Clara Moreau, Yassine Benhajali, Hien Duy Nguyen, Alan C. Evans, Pierre Bellec

**Affiliations:** Montreal Neurological Institute and Hospital, McGill University, 3801 Rue de l’Université, QC H3A 2B4, Montreal, Canada; Centre de Recherche de l’Institut Universitaire de Gériatrie de Montréal, 4565 Queen Mary Rd, QC H3W 1W5, Montreal, Canada; Centre de Recherche de l’Institut Universitaire en Santé Mentale de Montréal, 7401 Rue Hochelaga, QC H1N 3M5, Montreal, Canada; Département de Psychiatrie et d’Addictologie, Université de Montréal, Pavillon Roger-Gaudry, C.P. 6128, succursale Centre-ville, QC H3C 3J7, Montreal, Canada; Sainte Justine Research Center, University of Montreal, 3175 Chemin de la Côte-Sainte-Catherine, QC H3T 1C5, Montreal, Canada; Department of Mathematics and Statistics, La Trobe University, Plenty Rd & Kingsbury Dr, VIC 3086, Bundoora, Australia

## Abstract

Our understanding of the changes in functional brain organization in autism is hampered by the extensive heterogeneity that characterizes this neurodevelopmental disorder. Data driven clustering offers a straightforward way to decompose this heterogeneity into subtypes of distinguishable connectivity types and promises an unbiased framework to investigate behavioural symptoms and causative genetic factors. Yet the robustness and generalizability of these imaging subtypes is unknown. Here, we show that unsupervised functional connectivity subtypes are moderately associated with the clinical diagnosis of autism, and that these associations generalize to independent replication data. We found that subtypes identified robust patterns of functional connectivity, but that a discrete assignment of individuals to these subtypes was not supported by the data. Our results support the use of data driven subtyping as a data dimensionality reduction technique, rather than to establish clinical categories.

## Introduction

Autism spectrum disorder (ASD) is a prevalent neurodevelopmental condition of impaired social communication and restrictive behaviour, diagnosed in about 1% of children (***Lai et al., 2014***; ***Baio et al., 2018***), that is associated with extensive heterogeneity of behavioural symptoms and neurobiological endophenotypes (***Jacob et al., 2019***; ***Lombardo et al., 2019***). Functional magnetic resonance imaging (fMRI) has emerged as a promising technology to identify potential biomarkers of functional connectivity (FC) in ASD and other psychiatric disorders (***Castellanos et al., 2013***). However, efforts to characterize the functional brain organization in ASD have so far largely focused on case-control comparisons, thus ignoring the presumed heterogeneity of FC alterations (***Nunes et al., 2019***; ***Hahamy et al., 2015***).

Data driven cluster analysis has long been proposed as a solution to decompose the heterogeneity of behavioural symptoms in ASD into distinct subtypes (***Eaves et al., 1994***; ***Beglin****ge****r and Smith, 2001***), but these subtypes have proven difficult to distinguish in clinical practice (***Lord et al., 2012***) and were recently abandoned in favour of the broader concept of an autism spectrum (***American Psychiatric Association. and DSM-5 Task Force., 2013***). The lack of progress towards reproducible, brain based biomarkers of ASD (***Lombardo et al., 2019***) has renewed interest in clustering methods to decompose the heterogeneity of brain alterations into distinct subtypes that are hypothesized to underlie the multitude of behavioural symptoms.

To date, only a small number of studies have applied brain based subtyping to characterize the neurobiological heterogeneity in ASD and relate it to behavioural symptoms (***Hong et al., 2019***). Early work on subcortical volume alterations in ASD distinguished four subtypes, but did not find significant differences of behavioural symptoms between them (***Hrdlicka et al., 2005***). A more recent multi-modal analysis distinguished three subtypes of structural brain alterations in ASD and found that core ASD symptoms could be much better predicted from the structural MRI data when separate prediction models were trained on each subtype compared to the on the full, unstratified dataset (***Hong et al., 2017***). Work on the heterogeneity of FC in individuals with ASD, attention deficit hyperactivity disorder (ADHD), and NTC distinguished three FC subtypes among regions in the default mode network (DMN) and found that each subtype was associated with all three diagnostic groups, indicating that these FC subtypes may be shared across diagnostic boundaries (***Kernbach et al., 2018***). An analysis of whole-brain FC in ASD and neurotypical control (NTC) individuals distinguished two subtypes of diverging within- and between-network connectivity, but similarly showed that the assignment of individuals to these subtypes was not associated to their clinical diagnosis (***Easson et al., 2019***).

These initial findings of subtypes in ASD leave several important questions open. Firstly, studies have so far interpreted subtypes both as categories that individuals are discretely assigned to (***Hrdlicka et al., 2005***; ***Hong et al., 2017***) and as dimensions that each individual can have a continuous measure of similarity with (***Kernbach et al., 2018***; ***Easson et al., 2019***). However, the stability of either of these two methods of assigning individuals to subtypes has not been systematically established. Secondly, several previous studies have limited their investigation of subtypes to individuals who were already diagnosed with ASD (***Hrdlicka et al., 2005***; ***Hong et al., 2017***; ***Tang et al., 2019a***). Whether subtypes associated with ASD symptoms are specifically found among these diagnosed individuals, or are also prevalent in the general population has not been clearly established. Behavioural symptoms in ASD overlap with those of other neurodevelopmental disorders and also extend into the general population (***Constantino and Todd, 2003***; ***Grzadzinski et al., 2011***). Similarly, neurobiological endophenotypes associated with ASD have been shown to exist among individuals with other neuropsychiatric disorders (***Park et al., 2018***; ***Di Martino et al., 2013***). It is therefore important to investigate whether subtypes identified in mixed samples of both ASD and NTC individuals show an association with ASD diagnosis and symptoms. Thirdly, none of the brain based ASD subtypes reported in the literature have been replicated to date. The recent failure to replicate promising reports of clinically meaningful neuroimaging subtypes in depression (***Drysdale et al., 2017***; ***Dinga et al., 2019***) has highlighted the importance of this limitation for the autism literature.

In this work, we aim to address these three gaps by applying a straightforward, unsupervised subtyping approach to subdivide a heterogeneous sample of both ASD and NTC individuals by their network based FC patterns. Firstly, we systematically evaluate the robustness of the subtype maps, and the discrete and continuous assignment of individuals to them. Secondly, we determine whether diagnosis naive subtypes of FC show an association with clinical ASD diagnosis at the network level. And thirdly, we determine the generalizability of our findings by replicating them on an independent dataset.

## Results

### Subtype maps are stable

We first aimed at evaluating the robustness of subtype maps. Subtype maps are the spatial FC profiles corresponding to each identified subtype in the brain. For this purpose, we repeated the subtype analysis on random subsamples of 50% of the discovery dataset. We then matched the subtype maps of each seed network across subsamples, based on the highest similarity between pairs of maps. The average spatial Pearson correlation between matched subtype maps was 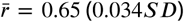 across all seed networks and subsamples. We observed small variations across seed networks: from *r* = 0.58 (0.081***SD***) for the inferior temporal gyrus seed network up to *r* = 0.7 (0.069***SD***) for the dorsal motor network (see ***Figure 1***). We thus showed that the subtype maps of the identified subtypes were robust to random perturbations in the dataset.

**Figure 1.**
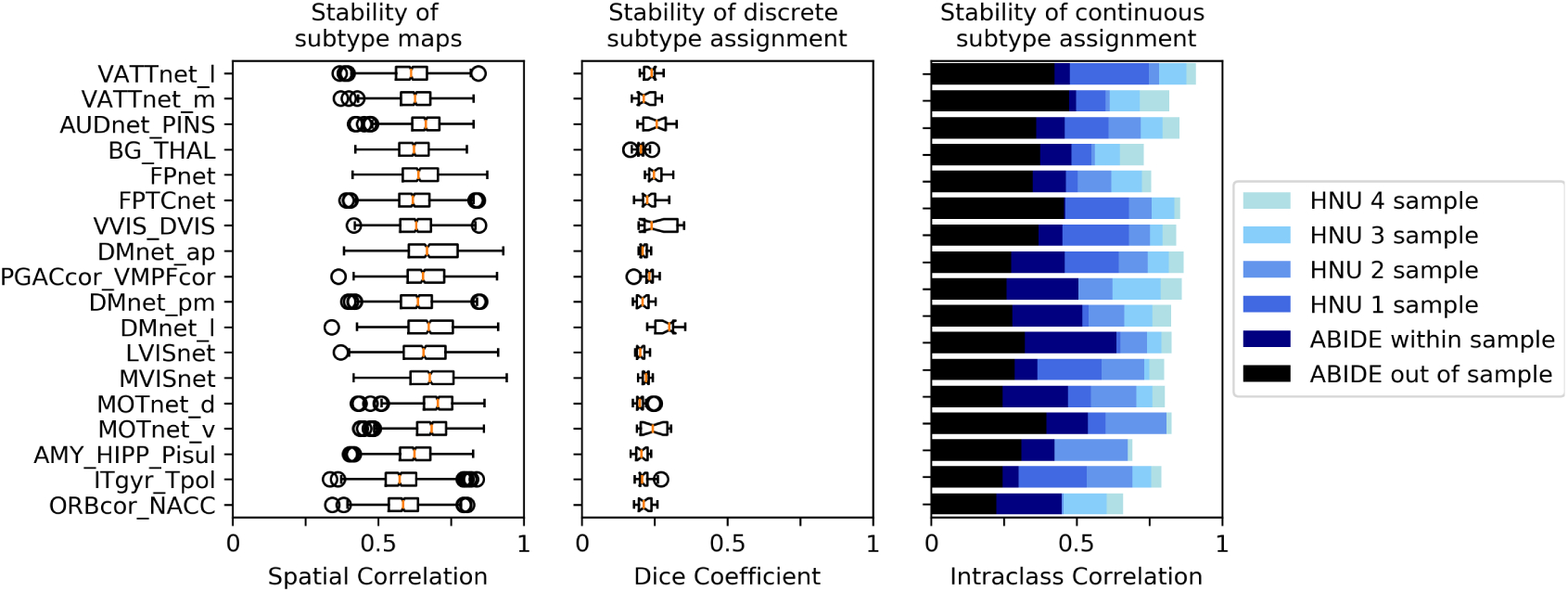
Robustness of subtyping outcomes across brain networks. Left: Stability of the FC subtype maps. Boxplots represent the range of the average similarity between FC subtype maps of the same brain network that were extracted from separate subsamples of the discovery dataset. Middle: Stability of discrete assignments of individuals to a FC subtype cluster. Boxplots represent the average overlap between the clusters an individual was assigned to in two different random subsamples. Right: Stability of continuous assignments of individuals to a FC subtype across repeated imaging sessions. Bar plots represent the average Intraclass Correlation between continuous subtype assignments computed on separate longitudinal imaging sessions. Different bar hues represent the stability of continuous subtype assignments extracted from out-of-sample subtypes (black), from within-sample subtypes (dark blue), within-sample subtypes in a general population data set where multiple scan sessions were combined to compute continuous subtype assignments (lighter shades of blue reflect more combined sessions).

### Discrete individual subtype assignments are not stable

To evaluate the robustness of discrete assignments of individuals to a subtype, we compared the overlap of the subtypes that an individual was part of between pairs of random subsamples. Because each subsample contained a random selection of individuals, we constrained our analysis to individuals that were included in both subsamples. The overlap was measured by the Dice coefficient (***Dice, 1945***). The average overlap of discrete subtype assignments was low at *Dice* = 0.22 (0.025***SD***). That is, 22% of the subtype neighbours of an individual in one subsample would on average also be subtype neighbours of this individual in another subsample. The range of overlap between subtypes was *Dice* = 0.2 (0.018***SD***) for the auditory network to *Dice* = 0.28 (0.046***SD***) for the medial visual network (see ***Figure 1***). We thus showed that the discrete assignment of an individual to a subtype was not robust to random permutations of the data in our simulation.

### Continuous individual subtype assignments are stable

We evaluated the replicability of continuous, individual subtype assignments for each seed network. To do so, we computed the intraclass correlation coefficient (ICC, ***Shrout and Fleiss, 1979***) of the continuous assignment across repeated scan sessions. The observed ICC coefficients were interpreted (***Cicchetti, 1994***) as

**poor** if less than 0.4

**fair** up to 0.59

**good** up to 0.74

**excellent** if larger than 0.75

When subtypes and the corresponding individual assignments to these subtypes were computed on data from the same individuals (within-sample replicability), the average ICC across seed networks was fair at ***ICC*** = 0.46 (0.073***SD***). The range of the stability of continuous subtype assignments across networks was ***ICC*** = 0.3 for the amygdala-hippocampal complex network to ***ICC*** = 0.63 for the lateral default mode network.

To evaluate the replicability of continuous subtype assignments, we repeated this analysis in a separate, general population dataset, wherein 10 scan sessions were available for each individual. In this data set, the average ICC of continuous subtype assignments was fair at ***ICC*** = 0.57 (0.094) when each assignment was computed on a single session. When estimating continuous assignments on the average of multiple sessions, the ICC increased markedly: good (***ICC*** = 0.68, ***SD*** = 0.09) for 2 sessions, good (***ICC*** = 0.75, ***SD*** = 0.071) for 3 sessions, and excellent (***ICC*** = 0.80, ***SD*** = 0.067) for 4 sessions.

Finally, we evaluated the replicability of continuous subtype assignments for subtypes that were computed on independent data (out of sample replicability). To this end, we computed subtypes on the discovery sample and estimated continuous subtype assignments for individuals in the longitudinal mixed patient-control sample. Here the average ICC was poor at ***ICC*** = 0.33 (***SD***0.072) with a range of ***ICC*** = 0.23 in the inferior temporal gyrus to ***ICC*** = 0.48 in the medial ventral attention network. We thus showed that the replicability of continuous subtype assignments ranges from poor to excellent as a function of the amount of available data per individual and whether subtypes and continuous subtype assignments are computed on the same data.

### Subtypes are robust to nuisance covariates and parameter changes

We then conducted the FC subtype analysis in the discovery dataset for each seed network. FC subtypes were identified according to two criteria: an average spatial dissimilarity below 1, and a minimum number of 20 individuals within each subtype. Across all seed networks, we identified 87 FC subtypes, with an average of 5 per network. These FC subtypes captured on average 97% of individuals in the sample (see also ***Appendix 1***). We tested whether continuous subtype assignments were driven by head motion, age, or recording site and found no significant linear associations with these covariates (see also ***Appendix 1***). Lastly, we evaluated whether our results were influenced by the choice of the dissimilarity threshold by repeating the subtyping and subsequent analysis steps for different levels of dissimilarity thresholds. We found that our results were robust across dissimilarity thresholds but that higher thresholds led to the inclusion of smaller proportions of the sample (see Fig 1 in ***Appendix 3***).

### Subtypes show association with ASD diagnosis

We next investigated whether any of the identified FC subtypes naturally captured interindividual variance of clinical ASD symptoms. To test this question, we computed the continuous assignment of all individuals in the discovery dataset to the identified subtypes. We then tested for a linear relationship between continuous subtype assignments and ASD diagnosis (i.e. ASD or NTC) and measures of ASD symptom severity (i.e. calibrated ADOS severity scores).

We identified 11 FC subtypes for which the continuous assignment of individuals were significantly associated with the clinical diagnosis of ASD, after correction for multiple comparisons (*p*_*adj*_ reflects the false discovery rate adjusted p-values, see Methods for details). That is, ASD and NTC individuals differed significantly in their continuous assignments with these subtypes. NTC individuals showed significantly stronger assignments than ASD individuals with 5 of the 11 subtypes. These protective subtypes originated from seed networks in the ventral motor network (*T* = 3.79, *p*_*adj*_ = 0.0037, *d* = −0.42), the auditory network (*T* = 4.25, *p*_*adj*_ = 0.0018, *d* = −0.52), the dorsal motor network (*T* = 4.15, *p*_*adj*_ = 0.0018, *d* = −0.49), the medial ventral attention network (*T* = 3.49, *p*_*adj*_ = 0.0091, *d* = −0.39), and the downstream visual network (*T* = 3.23, *p*_*adj*_ = 0.0196, *d* = −0.38).

ASD individuals showed significantly stronger continuous assignments than NTC individuals with 6 of the 11 subtypes. These risk subtypes originated from seed networks in the ventral motor network (*T* = 2.91, *p*_*adj*_ = 0.0330, *d* = 0.32), the dorsal motor network (*T* = 3.8−, *p*_*adj*_ = 0.0369, *d* = 0.39), the downstream visual network (*T* = 2.94, *p*_*adj*_ = 0.0330, *d* = 0.28), the amygdala-hippocampal complex (*T* = 2.75, *p*_*adj*_ = 0.0488, *d* = 0.27), the fronto-parietal control network (*T* = 2.92, *p*_*adj*_ = 0.0330, *d* = 0.29), and the lateral default mode network (*T* = 3.17, *p*_*adj*_ = 0.0204, *d* = 0.30). We thus showed that a subset of the identified FC subtypes naturally captured some variance of the clinical ASD diagnosis. We did not find an association between continuous subtype assignments and ASD symptom severity beyond the effect of the clinical diagnosis (see ***Appendix 2***).

### Subtype associations with ASD diagnosis replicate moderately

We next investigated how reproducible the discovered association between FC subtypes and ASD diagnosis was in an independent dataset. For each of the subtypes that showed a significant association with ASD diagnosis in the discovery dataset, we computed the continuous assignment for the individuals in the independent replication dataset. In this way, we tested the out of sample reproducibility of the observed association effect. We tested different degrees of replication: whether the observed effect in the replication sample was significant after correction for multiple comparisons, significant at an uncorrected *p* < 0.05, whether the estimated magnitude of the effect fell within the 90% confidence interval of the effect size estimate in the discovery sample, and whether the effect in the replication sample had the same direction as the one estimated in the discovery sample. We found that all effects of association with diagnosis in the replication sample had the same direction as in the discovery sample. The effect size estimates in the discovery sample were correlated at *r* = 0.91 with the effect size estimates in the replication sample. The magnitude of the estimated effects in the replication sample was on average 63% of those estimated in the discovery sample, and nine out of eleven effect size estimates fell within the 90% confidence intervals of the effect size estimates in the discovery sample. Five of those effects were significant at *p*< 0.05, and two of those were significant at *p*_*adj*_ < 0.05 (***Figure 2*** b). We thus showed that the association between subtypes and ASD diagnosis observed on the discovery dataset was moderately replicable in the independent replication dataset.

**Figure 2.**
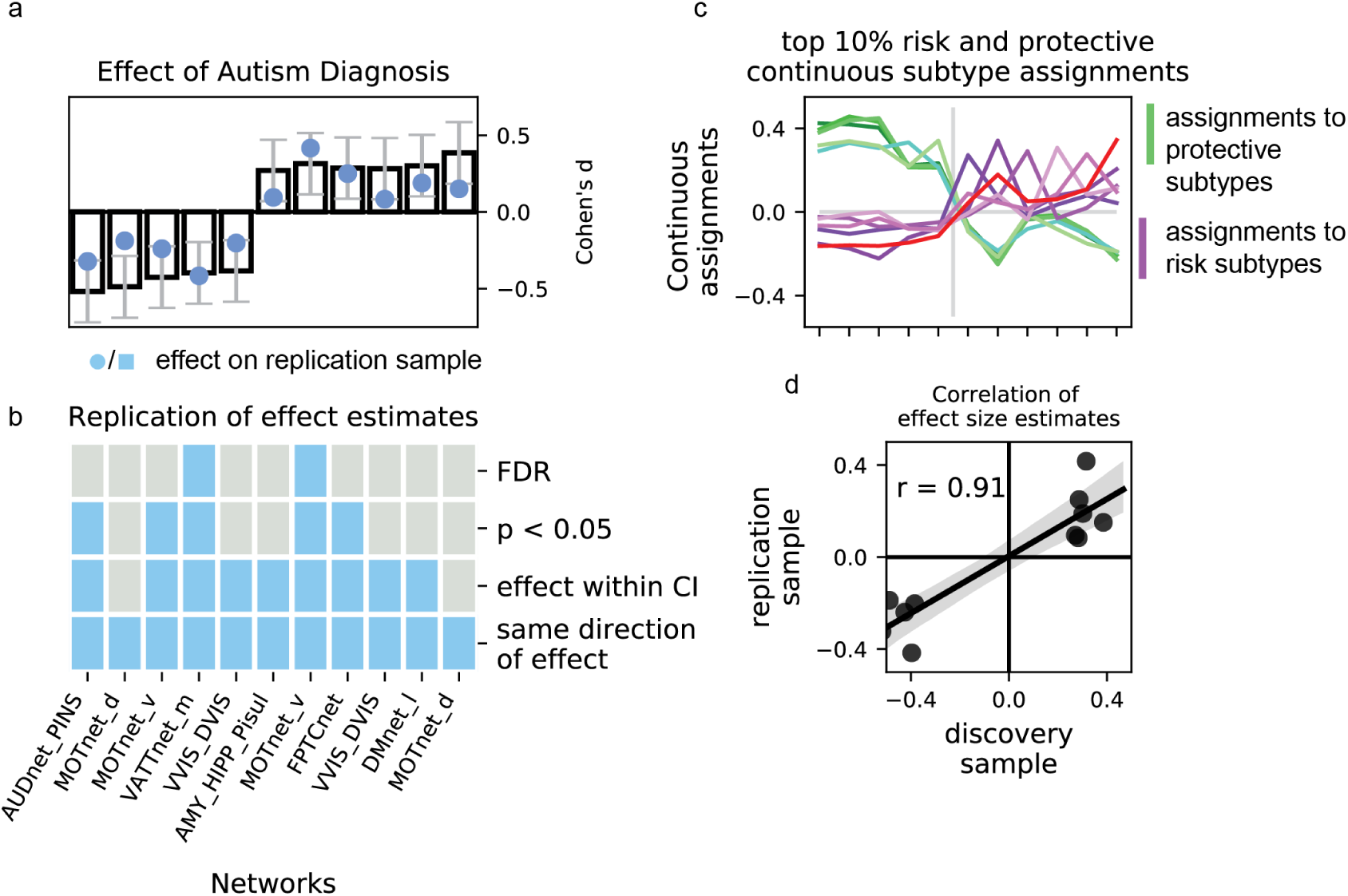
Association of continuous subtype assignments and diagnosis. a) Bar plots represent the standardized group difference (Cohen’s d) of continuous subtype assignments between NTC individuals and ASD patients. Negative value reflect greater similarity of neurotypical control subjects with the subtype, positive values reflect greater similarity of ASD patients with the subtype. Error bars reflect the 95% confidence interval of the effect size estimates. The effect size observed in the independent replication data set is shown as a blue dot. b) Matrix showing the degree of replication in the independent replication dataset of the observed association with diagnosis for each of the 11 protective and risk subtypes. Each row corresponds to a bar-plot in a). From top to bottom, the degrees of replication are: FDR: full replication of the effect after FDR correction, *p*< 0.05: replication of the effect for uncorrected statistics, effect within CI: observed effect size in the replication sample falls within the 95% confidence interval of the observed effect in the discovery sample, direction: observed effects in the discovery and independent replication sample go in the same direction. c) Graph illustrating the similarity of continuous subtype assignments across risk and protective subtypes. The average continuous subtype assignments of the top 10% of individuals with the highest similarity with a protective (green shades) or risk (red shades) subtype are displayed across all identified protective (left side) and risk (right side) subtypes. An individual may belong to the top 10% in more than one subtype. d) Correlation plot of the observed effect sizes in the discovery and independent replication datasets. The black line represents the correlation of effect sizes, the grey shaded area reflects the estimated 95% CI of the linear fit. **Figure 2–source data 1.** Table of the unthresholded association T-test between continuous subtype assignments and ASD diagnosis on the discovery dataset. **Figure 2–source data 2.** Table of the unthresholded association T-test between continuous subtype assignments and ASD diagnosis on the validation dataset.

### Subtypes with similar risk for ASD show similar spatial patterns of FC alterations

We noticed that the spatial pattern of protective subtype maps appeared similar, despite representing connectivity profiles from different seed networks (***Figure 3*** a, b). Similarly, the subtype maps of risk subtypes all appeared to show below average connectivity. We therefore investigated whether subtypes with the same direction of association with ASD diagnosis (i.e. protective and risk subtypes) shared similar FC profiles and whether this also extended to the continuous assignments of individuals to these subtypes. We found that protective subtypes exhibited a highly convergent pattern of FC alterations (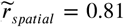, where 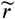 reflects the median spatial correlation across subtype pairs) that was distinct from those of risk subtypes 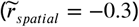. The spatial similarity among risk subtypes was less pronounced 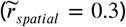 than that of protective subtypes. This finding extended to continuous subtype assignments that were more strongly correlated among protective subtypes 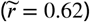 than among risk subtypes 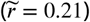, and anti-correlated between protective and risk subtypes 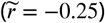. By dividing all subtype maps into the 18 seed networks, we observed that the shared spatial pattern of protective subtypes was characterized by overconnectivity with unimodal sensory brain networks, and underconnectivity with the basal ganglia and fronto-parietal network (green hues, ***Figure 3*** c). By contrast, the shared spatial pattern of risk subtypes was characterized by pervasive underconnectivity (red hues, ***Figure 3*** c). We thus showed that subtypes associated with a similar risk of ASD diagnosis exhibited similarities of FC alteration and continuous assignments, and that these similarities were more pronounced for protective subtypes than risk subtypes.

**Figure 3.**
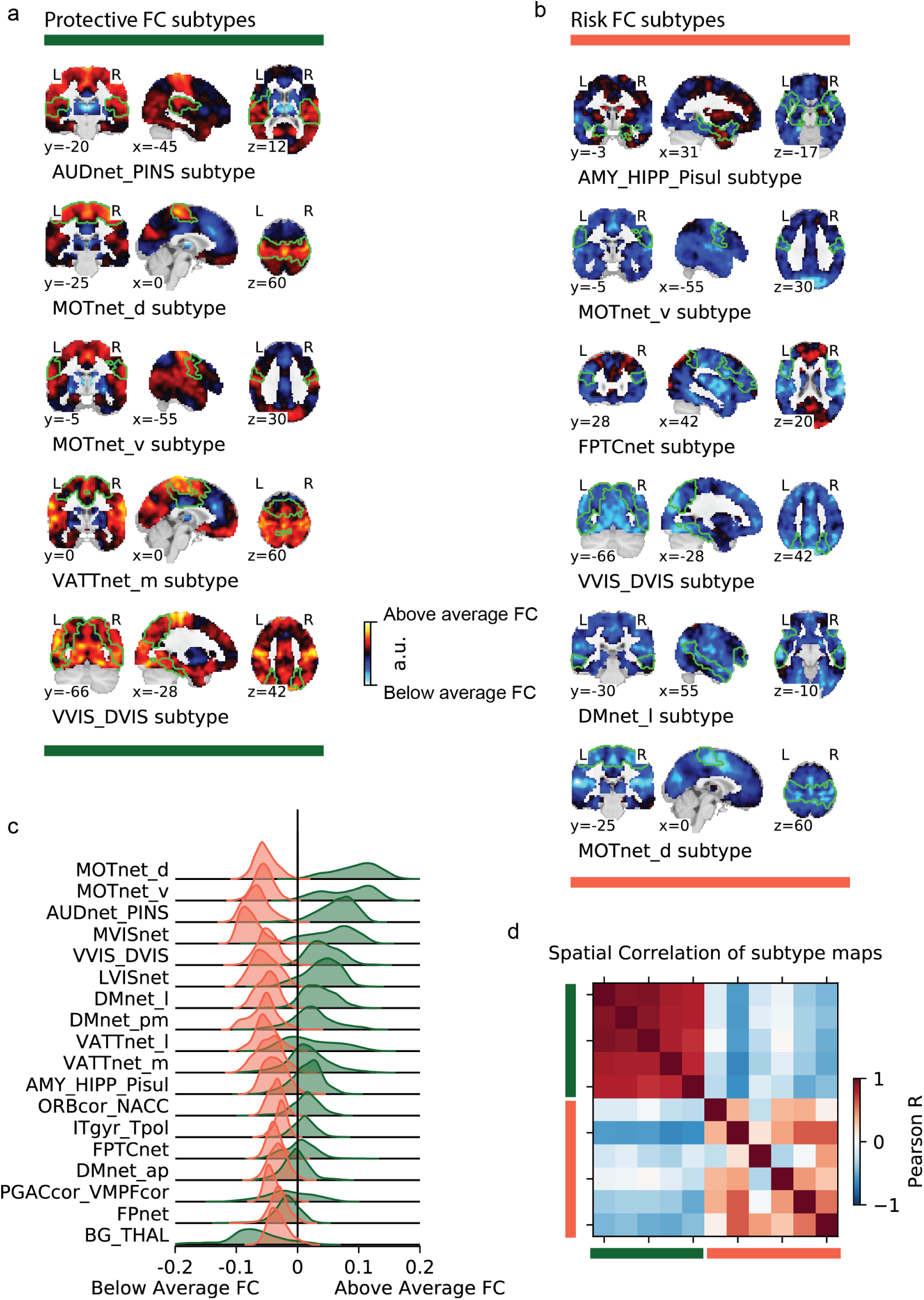
Overview of risk and protective subtype maps. Maps of protective (a) and risk (b) subtypes (corresponding seed networks are outlined with a thin green boundary on the map). c) Decomposition of the average protective (green) and risk (red) subtype map into 18 brain networks. d) Spatial correlation between subtypes. Protective (green) and risk (red) subtypes are denoted by colored bars along the correlation matrix.

## Discussion

ASD is characterized by a heterogeneity of symptoms and neurobiological endophenotypes (***Nunes et al., 2019***; ***Dickie et al., 2018***; ***Jacob et al., 2019***) among the affected individuals. Data driven, unsupervised subtyping appears as a natural approach to decompose the heterogeneity in ASD and to identify subtypes of functional brain connectivity. Here, we first sought to evaluate how stable and reproducible subtypes of FC are when derived from a heterogeneous sample of both neurotypical and autistic individuals. We then investigate whether fully data driven subtypes are associated with a clinical diagnosis of ASD. Our results suggest that data driven imaging subtypes are moderately reliable on currently available datasets and show a weak to moderate association with the clinical diagnosis of ASD, that generalizes to independent replication data.

### Functional connectivity subtypes are stable

Our systematic evaluation of the robustness of subtype maps, and the discrete or continuous assignments of individuals to them, establishes a foundation on which to understand previous incidental findings on the robustness (***Easson et al., 2019***) or non-reproducibility (***Dinga et al., 2019***) of subtype analyses. The FC patterns of the subtypes identified in our analysis were found to be robust to perturbations of the discovery data set. This observation fits with previous studies that reported stable imaging subtypes in ASD (***Hong et al., 2017***; ***Easson et al., 2019***; ***Tang et al., 2019b***). By contrast, we found that making a discrete assignment of individuals to the identified subtypes was not robust to perturbations. Few papers have investigated the robustness of discrete subtype assignments explicitly. One recent study attempted to replicate a high profile report of clinically predictive FC subtypes among depressed patients (***Drysdale et al., 2017***) but found that the asserted discrete subtypes were not sufficiently supported by an independent dataset (***Dinga et al., 2019***). The authors concluded that the data instead supported a more parsimonious model of continuous neurobiological axes.

In the wider ASD literature, the robustness of discrete subtype assignments has been more comprehensively investigated for symptom based subtypes. Several symptom based subtypes of autism have been proposed in attempts to provide more homogeneous diagnostic criteria. However, the distinction between these subtypes was also not found to be well supported by replication attempts which has led the field to merge sub-diagnoses of autism under the label of autism spectrum disorder (***Lord et al., 2012***; ***Volkmar and McPartland, 2014***).

We may reconcile the seemingly conflicting findings of robust subtypes on the one hand and non-reproducible discrete subtype assignments on the other, when we consider that the similarity of an individual with each subtype is a continuous measure. For individuals who are equally similar to two different subtypes, a small change of the connectivity profile of either of the subtypes may be enough for them to be assigned to the other subtype if a discrete choice is forced. By contrast, the continuous similarity measure would not change drastically. An emerging body of literature therefore conceptualizes subtypes as latent dimensions that can be expressed to varying degrees in each individual (***Kernbach et al., 2018***; ***Tang et al., 2019b***; ***Easson et al., 2019***). Our own results support this view: we find that unlike discrete subtype assignments, continuous measures of an individuals’ similarity with each subtype are moderately robust and can be very robust when more data is available per individual to compute the continuous assignment. The ICC of continuous subtypes assignments computed on separate data was low but consistent with previous reports of the robustness of single session seed based FC measures (***Shehzad et al., 2009***). When the continuous subtype assignments were computed based on the average FC of multiple scan sessions per individual, we found high to very high robustness measures that were in line with the well established link between scan length and FC reliability (***Gordon et al., 2017***). It is reasonable to assume that the generalizability of the associations between continuous subtype assignments and clinical ASD diagnosis we have reported here could be increased if longer or repeated scan sessions were available for the replication sample. Based on our findings we may thus conclude that continuous measures of individual subtype assignment that reflect the similarity of an individual with several subtypes provide a better representation of the data.

### Subtypes moderately, but reproducibly, associate with ASD diagnosis

The majority of previous subtyping analyses in ASD have been constrained to patients that were already diagnosed with ASD (***Hrdlicka et al., 2005***; ***Hong et al., 2017***; ***Tang et al., 2019b***). We have instead used an unsupervised clustering approach to identify diagnosis-naive subtypes of FC across autistic and neurotypical individuals in order to determine whether they would associate with ASD diagnosis. Our results showed that these subtypes were significantly associated with a clinical diagnosis of ASD, and that the observed effects were small to moderate, ranging between *d* = 0.3 and *d* = 0.5 (or from *r* = 0.15 to *r* = 0.24 when expressed as a correlation coefficient) on the discovery sample, with reduced effect sizes identified in an independent replication sample.

Our effect sizes are comparable to those reported by other imaging based subtypes in ASD, which have all estimated association with diagnosis in their discovery sample (i.e. have not been replicated on independent data). A recent study (***Kernbach et al., 2018***) investigated the heterogeneity of FC in mixed data of ASD, NTC, and attention deficit hyperactivity disorder (ADHD), a common comorbidity of ASD individuals (***Rommelse et al., 2010***). The authors identified one FC endophenotype that was weakly associated with ASD (*r* = 0.15) but extended both to ADHD and NTC individuals. Another study on structural imaging subtypes among ASD (***Hong et al., 2017***) patients found that ADOS severity scores could be better predicted from structural cortical alterations when individuals were first divided into three subtypes (*r* = 0.47 compared to *r* = −0.12 when the association was computed without regard for subtypes), although the prediction performed worse for calibrated ADOS severity (*r* = 0.25). The magnitude of the association between data driven subtypes and clinical diagnosis in our analyses is therefore comparable to what has been previously reported by other imaging based subtypes analyses of ASD. Weak-to-moderate associations between data driven subtypes and clinical diagnosis thus seem robust to the employed subtyping method, at least in the currently limited number of published studies.

Very few imaging based subtype analyses have been replicated on independent data, and to our knowledge none have so far been replicated successfully (***Dinga et al., 2019***). The replication of our results on independent data therefore establishes a novel benchmark of reliability for imaging based subtype analyses in ASD. We found that the observed effect sizes of the association between FC subtypes and clinical ASD diagnosis strongly correlated between the discovery dataset and the independent replication dataset (*r* = 0.91), however effect sizes in the replication data were on average only 2/3rds the the magnitude of those in the discovery data. This reduction of effect sizes on the replication data is expected as it reflects the inherent bias of significance testing to select larger effects and further underlines the importance of reporting original findings together with independent replications for an unbiased estimate (***Vul and Pashler, 2012***). Because no other imaging subtype analysis in ASD has been independently replicated to date, our results have to be interpreted in the context of replication attempts in the ASD case-control literature. The largest case-control analysis of FC alterations to date (***Holiga et al., 2019***) reported FC group differences between ASD and NTC individuals with effect sizes between *d* = 0.46 and *d* = 0.6, similar in size to our own results of FC subtype associations with clinical ASD diagnosis. Using several large replication samples, the authors then showed that these results were reproducible in independent data, however with similarly depressed effect sizes (i.e. *d* ≈ 0.2).

The comparison with the large case-control study by Holiga and colleagues may also serve to illustrate the conceptual advantage of a subtyping approach over the traditional case-control design for the investigation of ASD related alterations of FC. An incidental finding of our analysis was that protective subtypes converged onto a network of mutually overconnected and predominantly unimodal brain networks (see ***Figure 3***). The spatial pattern of this overconnectivity profile is visually very similar to the case-control pattern of differences FC ASD and NTC individuals reported in the study of Holiga et al (see ***Figure 4***). We computed the spatial correlation between the two patterns at *r* = −0.6, as high or higher than the reported replicability of the case-control pattern itself (between *r* = 0.3 and *r* = 0.6, depending on the replication sample). Because the case-control results of Holiga are at least in part based on the same data that were used in our study, the strikingly high spatial correlation and the similarity of effect sizes suggests that case-control studies may not capture ASD specific FC alteration patterns but are rather driven by the most prevalent FC subtypes in the data, that are themselves not strongly linked to ASD. The fact that our analysis identified several distinct and non-overlapping FC subtypes that were reproducibly associated with moderately increased risk of ASD further illustrates the conceptual limitation of computing group averages across data known to be highly heterogeneous (***Ecker and Murphy, 2014***; ***Lombardo et al., 2019***).

**Figure 4.**
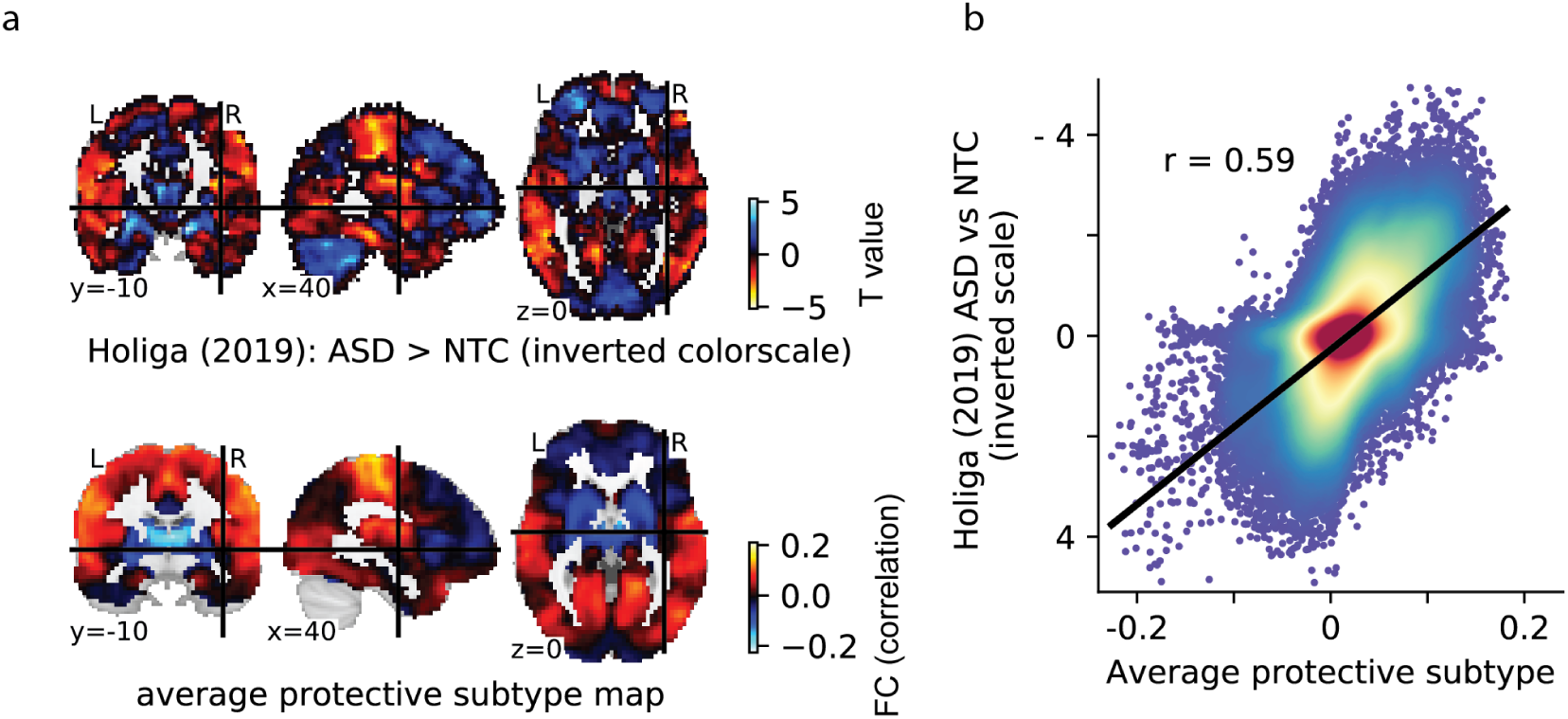
Comparison of the average protective subtype map to a case-control signature. a) The spatial map of a large sample size case-control contrast between ASD and NTC individuals (top row), compared to the average spatial map of the protective subtypes identified on our data (bottom row). Note that because of the opposite nature of the two contrasts (i.e. ASD > NTC for the case-control contrast and NTC > ASD for the protective subtype map), the color scale for the case-control map has been inverted for better comparability. b) Plot of the voxel-wise spatial correlation between the (inverted) case-control contrast map and the average protective subtype map. The blue to red color gradient reflects the density of voxels represented in each area of the graph.

Taken together, our results suggest that unsupervised FC subtypes associate with clinical ASD diagnoses at a level similar to case-control studies and have comparable reproducibility. They also likely provide a more comprehensive and informative representation of the underlying neurobiological heterogeneity.

### Limitations

Our findings are limited by the amount of data available per individual. We found that continuous assignments of individuals to subtypes became more stable, the more data was available per individual. One reason for this observation could be that the FC pattern associated with an individual is in fact not static across time but rather represents an average of a number of dynamic states (***Allen et al., 2014***). How long an individual spends in each dynamic state can be a reproducible trait (***Choe et al., 2017***) and may provide additional insight into the relationship of connectivity and ASD symptoms (***Rashid et al., 2018***). It is therefore possible that by averaging longer time series for each individual, we get a better approximation of that individual’s preferred dynamic state. A promising direction for future research will be the investigation of dynamic FC subtypes in ASD. Datasets providing longer time series per individual will facilitate these inquiries.

Our results have focused only on individuals with ASD. Given the extensive evidence of overlap of symptoms (***Grzadzinski et al., 2011***) and neurobiological phenotypes between ASD and other neurodevelopmental disorders (***Sha et al., 2019***), a fruitful avenue for future research will be to extend this approach to investigate cross-diagnostic subtypes of FC (***Elliott et al., 2018***).

### Conclusions

Our findings suggest that unsupervised clustering of heterogeneous imaging data is well suited to identify subtypes that are reproducibly associated with clinical symptoms. The low to moderate effect size of the observed association makes it clear that subtypes of FC will not replace or provide better clinical categories than the current diagnostic system. The instability of discrete assignments of individuals to subtypes and the small to moderate effect sizes associated with these subtypes do not lend themselves to make meaningful predictions about the clinical prognosis of an individual. However, we find that the associations between subtypes and clinical diagnosis do generalize to independent data and that the risk subtypes we identified in our dataset capture non-redundant profiles of FC dysfunction. Both of these observations point towards a promising avenue for future research: data driven subtyping appears well suited as an efficient method to summarize high dimensional, heterogeneous data while retaining clinically meaningful variation. These properties make FC subtypes good candidates as features for multivariate, supervised predictive learning models to predict clinical diagnosis.

## Methods and Materials

### Discovery sample

The discovery sample consisted of imaging data from the ABIDE 1 dataset (***N*** = 388, ***N***_*A****SD***_ = 194, ***A****ge* = 17.04, (7.08), from 7 recording sites) and ASD individuals were matched with neurotypical controls on age (***A****ge*_*A****SD***_ = 17.0, (7.28); ***A****ge*_*NTC*_ = 17.04, (6.89)) and head motion (***FD***_***ASD***_ = 0.17*mm*, (0.048); ***FD***_*NTC*_ = 0.16*mm*, (0.041)). The full ABIDE 1 dataset includes 1112 individuals from 20 imaging sites (***N***_***ASD***_ = 539, ***a****ge* = 17.04, (8.04)) of which 948 are male. Due to the strong sex imbalance of the data, we limited our analysis to male individuals. After preprocessing of the imaging data, 557 individuals (272***ASD, A****ge* = 16.65, (6.75)) from 13 imaging sites were found to pass our quality control criteria. We then matched the NTC and ASD individuals at each site by age and head motion through propensity score matching without replacement (***Rosenbaum and Rubin, 1985***). The matched sample included 478 individuals (239***ASD, A****ge* = 16.67, (6.67)). We further excluded the 5 imaging sites with fewer than 20 matched individuals, leaving 388 individuals for the final discovery sample.

### Replication sample

The replication sample consisted of imaging data from the ABIDE 2 dataset (***N*** = 300, ***N***_*A****SD***_ = 150, from 7 imaging sites) and ASD individuals were matched with neurotypical controls on age (***A****ge*_*A****SD***_ = 12.0, (4.05); ***A****ge*_*NTC*_ = 12.3, (4.59)) and head motion (***FD***_*A****SD***_ = 0.17, (0.053); ***FD***_*NTC*_ = 0.16, (0.048)). The full ABIDE 2 dataset includes 1114 individuals from 19 imaging sites (***N***_*A****SD***_ = 521, ***A****ge* = 14.86, (9.16)) of which 856 are male. Analogous to the discovery sample we limited our analysis to male subjects. After preprocessing of the imaging data, 587 individuals (***N***_*A****SD***_ = 273, ***A****ge* = 13.94, (5.9)) from 16 imaging sites were found to pass our quality control criteria. These individuals were then matched by age and head motion within each site through propensity score matching without replacement. The matched sample included 424 individuals (***N***_*A****SD***_ = 212, ***A****ge* = 13.66, (5.25)). We further excluded 9 imaging sites with fewer than 20 matched individuals, leaving 300 individuals for the final replication sample.

### Longitudinal sample 1

The first longitudinal test sample was taken from a subset of individuals in ABIDE 2 for whom multiple scan sessions were available. In ABIDE 2, longitudinal imaging data are available for 168 individuals from 4 imaging sites (***N***_*A****SD***_ = 88, ***A****ge* = 21.24, (15.45)) of which 154 are male. Analogous to the discovery and replication sample, we limited our analysis to male individuals. After preprocessing of the imaging data, 84 individuals (***N***_*A****SD***_ = 42, ***A****ge* = 14.58, (6.29)) from 3 imaging sites were found to pass our quality control criteria. We selected the two imaging sites with the largest number of acceptable individuals (ABIDEII-OHSU_1 and ABIDEII-IP_1) and randomly selected individuals at each site to enforce equal sized groups of NTC and ASD. Where more than 2 acceptable imaging scans were available for an individual, the 2 scans with the lowest average head motion were selected. The final longitudinal test sample consisted of 68 individuals (***N***_*A****SD***_ = 34, ***A****ge* = 13.46, (5.79)) from 2 imaging sites.

### Longitudinal sample 2

The second longitudinal test sample consisted of individuals in the general population Hangzhou Normal University dataset (http://dx.doi.org/10.15387/fcp_indi.corr.hnu1) released by the consortium for reliability and reproducibility (***Zuo et al., 2014***). The final sample included 26 individuals (***N***_*male*_ = 14, ***A****ge* = 24.58, (2.45)) that were each scanned 10 times at 3 day intervals over the course of a month. We selected the 26 individuals (out of a total of 30 available individuals) for which all resting state scans passed visual quality control.

### Clinical scores and symptom severity

The individuals from the ABIDE 1 and ABIDE 2 samples included in this study were diagnosed with ASD by expert clinicians based on either the Autism Diagnostic Observation Schedule (ADOS) (***Lord et al., 2000***; ***Gotham et al., 2007***) or the Autism Diagnostic Interview - Revised (***Lord et al., 1994***). The ADOS provides a total sum of ratings of observation items in the ADOS subdomains that reflects the severity of observed symptoms but is primarily intended for diagnostic purposes. Calibrated ADOS severity scores have been proposed as a standardized, research appropriate measure of symptom severity that is comparable across ADOS modules and is less dependent on demographic factors such as age (***Gotham et al., 2009***). Few individuals in the discovery (N=109, 93 ASD) and replication (88 ASD) samples had calibrated ADOS severity scores. We therefore also investigated associations with the raw ADOS total scores that were available in larger numbers in the discovery (***N*** = 213, ***N***_*A****SD***_ = 182) and replication (***N*** = 157, ***N***_*A****SD***_ = 148) sample.

### Imaging data preprocessing

All imaging data were preprocessed with the NeuroImaging Analysis Kit (NIAK) version 1.13 (***Bellec et al., 2011***). The preprocessing pipeline was executed inside a Singularity (version 2.6.1) software container (***Kurtzer et al., 2017***) to facilitate the reproducibility of our findings. Preprocessing of the functional imaging data consisted of the following steps: Head motion between frames was corrected by affine realignment with a reference image (median image across frames). The magnitude of framewise head displacement (FD) was estimated from the time course of the affine realignment parameters (***Power et al., 2012***). The reference image was then coregistered into the MNI152 stereotaxic space (***Evans et al., 1994***) through an initial affine alignment with the individual anatomical T1 image and a subsequent, non-linear coregistration of the T1 image with the MNI template. A high-pass temporal filter (0.01 Hz) was fitted to the whole time series by discrete cosine transform to remove slow time drifts. Time frames with excessive head motion (***FD*** > 0.4*mm*) were then censored by removing the affected frame, as well as the preceding and the two succeeding frames from the time series (***Power et al., 2012***). Nuisance covariates were then regressed from the remaining time points: the previously estimated discrete cosine basis functions, the average signal in conservative masks of the white matter and lateral ventricles, and the first principal components (accounting for 95% of variance) of the six rigid-body motion parameters and their squares (***Lund et al., 2006***; ***Giove et al., 2009***).

### Quality control of imaging data

We controlled the quality of preprocessed data manually and through quantitative cut off values. Data were visually checked by a trained rater following a standardized QC protocol (***Benhajali et al., 2020***) with a structured QC tool (***Urchs et al., 2018***). Individuals were excluded for coregistration failure and for incomplete brain coverage of the field of view of the functional data. During the visual QC we noticed that a large number of individuals in both the discovery and replication sample had incomplete field of view coverage of the cerebellum. We chose to remove all cerebellar networks from our analyses in order to include these individuals. Individuals were also excluded from the analysis if fewer than 50 time frames remained after motion censoring or if the average framewise displacement exceeded 0.3 mm.

### Functional connectivity estimation

We estimated the seed based FC maps of 18 non-cerebellar seed networks defined in the MIST_20 functional brain atlas (***Urchs et al., 2017***). The MIST_20 atlas represents large, spatially distributed subcomponents of canonical FC networks. The seed to voxel FC maps were estimated as the Pearson correlation between the average time series signal of a seed network and the time series of all gray matter voxels in the brain (excluding the cerebellum). Within each sample separately, the individual seed FC maps were centered to the group mean and known sources of variance of non-interest were regressed for each voxel at the group level: linear effects of age, head motion and imaging site. As a consequence, the individual seed FC maps in each sample represented the residual variance around the group mean after accounting for these factors.

### Subtyping of functional connectivity

To identify communities of individuals with similar seed FC patterns we computed the spatial correlation of all pairs of subjects in the discovery sample, separately for each seed network. We expressed the dissimilarity between pairs of individual seed FC maps as the absolute value of 1 - their spatial correlation. The 18 subject by subject dissimilarity matrices (one per seed network) thus contained values between 0 (no dissimilarity or a spatial correlation of 1) to 2 (perfect dissimilarity or a spatial correlation of −1) with 1 denoting no spatial relationship (a spatial correlation of 0).

For each seed network separately, we characterized communities of individuals with similar seed FC maps by hierarchical agglomerative clustering of the dissimilarity matrix for each seed network using the unweighted average distance linkage criterion (***Müllner, 2011***). We applied two criteria for the identification of seed FC communities: 1) the average dissimilarity between seed FC maps in a community could not be greater than 1, and 2) the community had to have at least 20 members. This allowed for small subsets of individuals with distinct seed FC patterns to not be assigned to any communities. Assigning individuals to subtypes in this way is a discrete process and we therefore refer to these assignments as discrete subtype assignments.

Within each seed FC community, we estimated the average seed FC map across all community members. This map reflected the subtype of seed FC shared by the community members and we refer to these maps as the subtype map.

Finally, we computed the spatial similarity of each individual in the discovery sample with the identified seed FC subtypes by spatial correlation of the individual seed FC map with the corresponding seed FC subtype map. The estimated spatial correlation coefficient is a continuous measure of an individual’s similarity with each of the subtypes and we therefore refer to it as a continuous subtype assignment. Each individual had continuous subtype assignments for each identified subtype, ranging from −1 (perfect anticorrelation of the individual and the subtype seed FC map) to +1 (perfect correlation of the individual and subtype seed FC map).

### Stability analysis

Before we investigated the three aspects of FC subtypes (subtype maps, and discrete and continuous assignments) in detail, we wanted to determine the robustness of these metrics to perturbations of the discovery data. We used two approaches: 1) to determine the robustness of discrete subtype assignments and subtype maps, we conducted a stratified subsampling scheme on our discovery sample, 2) to determine the robustness of continuous subtype assignments, we computed the within subject stability of continuous subtype assignments across repeated scan sessions for individuals in the longitudinal sample.

We randomly selected 1000 stratified subsamples of half of our discovery sample while preserving the equal ratio of ASD patients and NTC. Within each subsample, we repeated the full subtype characterization procedure: group level regression of nuisance sources of variance, characterization of communities of similar residual seed FC maps, estimation of seed FC subtype maps. The number of unique pairs of subsamples was large (≈ 500.000) and there was considerable overlap of individuals between subsamples. Therefore, we randomly selected 1000 unique pairs of sub-samples to estimate the robustness of the subtype community membership and subtype maps to perturbations in the data.

We determined the robustness of discrete subtype assignments by computing the similarity of the communities an individual was assigned to within two subsamples using the Dice coefficient (***Dice, 1945***). For each pair of subsamples A and B, we first identified the intersect of individuals (i.e. those individuals that were present in both subsamples). For each individual we then computed the Dice coefficient of the communities it was assigned to in sample A and sample B. The Dice coefficient here computes the ratio of twice the number of individuals shared between both communities over the total number of individuals in both communities. Thus, if all community neighbours of an individual in sample A were also community neighbors of that individual in sample B, then the Dice coefficient will be 1. Conversely, if none of the community neighbours of an individual in sample A were community members of that individual in sample B, then the Dice coefficient will be 0. We computed the average Dice coefficient across all individuals shared between a pair of subsamples.

We determined the robustness of the subtype maps by examining the spatial correlation of subtype maps extracted in each pair of subsamples. For each pair of subsamples A and B, we computed the spatial correlation of all subtype maps in sample A with all subtype maps in sample B. If subtype maps were robustly identified, then we would expect that for each subtype map in sample A we can find at least one subtype map in sample B that is very similar. We therefore searched (with replacement) for each subtype map in sample A the subtype map in sample B with the highest spatial correlation. Since the number of subtypes extracted in each subsample was determined by the data, we allowed for subtype maps in sample B to be a match for multiple subtype maps in sample A. We then took the average of the maximal spatial similarity between subtype maps of sample A and B as a measure of the robustness of the subtype maps.

We computed the robustness of the continuous subtype assignments as the intraclass correlation coefficient between repeated scan sessions of the same individual. We first investigated the robustness of assignments to subtypes that had been identified on data from a separate scan session but of the same sample (within sample robustness). Using the longitudinal sample 1, we identified FC subtypes for each network on scan session 1, and computed seed based FC maps for all individuals on the remaining two scan sessions. Independently for each scanning session we then centered the seed FC maps to the group mean and regressed covariates of non-interest for each voxel. The residual seed FC maps were then used to compute the continuous subtype assignments for the FC subtypes identified on scan session 1. The replicability of these continuous subtype assignments across the two remaining scan sessions was then estimated with the intraclass correlation coefficient.

Using the longitudinal sample 2, we tested whether continuous subtype assignments were more robust if they were computed on larger amounts of data per individual. Again, we identified FC subtypes for each network on the first scan session and computed individual seed FC maps on the remaining nine scan sessions. Within each scan session, the individual seed FC maps were then centered to the group mean and nuisance covariates were regressed. Two average residual seed FC maps per individual were then computed by averaging across sets of 2, 3, and 4 scan sessions. We then computed the continuous subtype assignments with the FC subtypes and estimated their replicability for the different number of averaged scan sessions with the intraclass correlation coefficient.

Finally, we computed the out of sample robustness of continuous subtype assignments based on the repeated scan session in the longitudinal sample 1 with the FC subtypes identified on the complete discovery sample. Again, the robustness was measured with the intraclass correlation coefficient of continuous subtype assignments across scan sessions.

### Association with autism diagnosis

We explored whether seed FC subtypes existed for which the presence of an autism diagnosis explained a significant amount of variance of the continuous subtype assignments. We tested this for each subtype by comparing the means of continuous subtype assignments between ASD individuals and NTC with a general linear model with diagnosis as the explanatory factor. As we had taken care to ensure equal sizes of individuals in both diagnostic categories, we did not use a correction for unequal variances. The estimated p-values were corrected at a false discovery rate (FDR) of 5% across all subtypes using the Benjamini and Hochberg method (***Benjamini and Hochberg, 1995***). We report the standardized group difference (Cohen’s d) between diagnosis and continuous subtype assignments as a measure of the effect size of the association with the clinical diagnosis. We continued investigating subtypes for which a significant difference of continuous subtype assignments between ASD patients and NTC was found in the discovery sample.

Within the set of subtypes that showed a significant association with ASD diagnosis we investigated whether spatial similarity with the subtype map explained additional variance of the severity in clinical symptoms. Because symptom severity and the clinical ASD diagnosis were highly correlated, and because healthy individuals had compressed or missing scores for most severity measures, we only tested this association in individuals with a diagnosis of ASD. We investigated the linear relationship between continuous subtype assignments and severity estimates for the calibrated ADOS severity scores (***Gotham et al., 2009***) and also for the raw ADOS total scores. We reported the correlation between symptom scores and continuous subtype assignments as a measure of the effect size of the association with symptom severity after correction for multiple comparisons using FDR.

### Replicability

We tested the replicability of the associations between seed FC subtypes and ASD diagnosis in an independent replication sample. Within the replication sample we computed individual seed FC maps for the 18 non-cerebellar MIST_20 seed networks, centered the seed FC maps to the replication sample group average and regressed variance of non-interest due to age, head motion and imaging site for each voxel. For the residual seed FC maps, we computed the continuous subtype assignment scores with the subtypes identified in the discovery sample. For those subtypes that showed significant associations with ASD diagnosis in the discovery sample, we then investigated the difference in continuous subtype assignment scores between ASD and NTC individuals in the replication sample.

### Robustness of findings to changes in the subtyping pipeline

Although we did not explicitly specify the number of subtypes to be identified for each seed network, it was implicitly determined by the maximum dissimilarity parameter and the structure of the subject by subject dissimilarity matrix. In order to understand how robust our findings were to changes in this parameter, we repeated all analysis steps (i.e. the identification of subtypes, the test for associations with ASD symptoms, and the generalization to the independent replication data) for different values of the maximum dissimilarity parameter. To measure the spatial similarity of subtype maps identified for different dissimilarity parameters we computed their pairwise spatial correlation. We then compared the number of identified subtypes and the observed associations with ASD symptoms and their generalization to independent data qualitatively.

## Acknowledgments

This research was supported by computation resources of Calcul Quebec and Compute Canada. We thank Gleb Bezgin, Budhachandra Khundrakpam, Yasser Iturria Medina, and John Lewis for helpful discussions of the analytic concept. For their feedback on the writing of this manuscript we want to thank Julie Boyle, Nida Ali, and Jonas Nitschke. We thank the ABIDE consortium and the consortium for reliability and reproducibility (CORR) for making publicly available the large datasets that this study was based on.

## Appendix 1

### Subtypes capture majority of individuals in the dataset

Subtypes for each seed network were identified according to two criteria: the average spatial dissimilarity within a subtype was no larger than 1 and at least 20 individuals were part of the subtype. Across all 18 seed networks, we identified 87 FC subtypes in the discovery dataset. In each seed network we identified between 3 (medial visual network) and 6 sub-types (lateral visual network) that satisfied these criteria (the median number of subtypes was 5). On average across networks, 97% of the individuals in the discovery dataset were assigned to a subtype (see also Figure 5). The largest number of individuals not assigned to any subtype was 19 in the inferior temporal gyrus network, and all individuals were assigned to subtypes in the ventral somatomotor and perigenual anterior cingulate seed networks. The average number of individuals in a subtype was 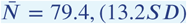. We thus show that the majority of individuals in the discovery dataset contributed to the identified 87 FC subtypes.

### Subtypes are not driven by nuisance covariance

To ensure that subtypes were not driven by variation of non interest, we tested for linear associations between the continuous assignment of individuals to each subtype, and head motion and age by Pearson correlation. We also tested whether recording sites were over-represented in subtypes above chance level with a chi-square test. We found no significant linear relationship between continuous assignments of individuals to subtypes and in-scanner head motion, and age for in any seed network. In addition, we found that the distribution of imaging sites across subtypes did not differ significantly from chance. We thus show that subtypes in the discovery dataset were not significantly driven by variance sources of non-interest.

## Appendix 2

### No added effect of ASD symptom severity beyond diagnosis

We also investigated whether any of the identified subtypes captured variation due to ASD symptom severity beyond the observed effects of ASD diagnosis. The distinction between the effects of ASD diagnosis and additional effects of ASD symptom severity is necessary because ASD symptom severity measures are by definition strongly correlated with ASD diagnosis. Calibrated ADOS symptom severity scores were available for 109 individuals from four recording sites in the discovery dataset (NYU, UCLA, KKI, USM). Of those, only 16 were NTC. In the independent replication dataset, calibrated ADOS severity scores were available for 88 individuals from five recording sites (NYU_1, OHSU_1, SDSU_1, KKI_1, GU_1). All of the 88 individuals were ASD individuals. We therefore limited our analysis to ASD patients. When controlling for the effect of clinical diagnosis in this way, we did not find significant additional effects of symptom severity in any subtypes. Repeating this analysis for the ADOS raw total scores (that were available for 182 and 148 ASD individuals in the discovery and replication sample respectively) likewise resulted in no signi_cant association with continuous subtype assignments for the identified risk and protective subtypes. We thus showed that beyond the effects of clinical ASD diagnosis, there was no significant added effect of symptom severity captured by the FC subtypes.

## Appendix 3

### Effects are robust to changes in the subtyping method

We identified subtypes of FC that satisfied two criteria: a maximal average dissimilarity of the connectivity patterns of individuals contributing to the subtype, and a minimal number of individuals. Although this process did not explicitly specify the number of subtypes to be identified, we sought to understand how robust our findings were to changes in the subtyping criteria. We therefore repeated the complete subtype analysis (i.e. identification of subtypes, association with ASD diagnosis, and generalization on independent data) for different values of maximal within-subtype dissimilarity. This analysis revealed that subtype maps remained highly similar across different values of the dissimilarity criterion with subtypes for the most part contracting and only rarely splitting into subcomponents (see Figure 4). We found highly consistent spatial patterns of protective and risk subtypes and effects of association with clinical ASD diagnosis did generalize to the independent replication dataset at equal rates. We thus conclude that our findings were robust to changes in the parameters of the subtyping analysis.

**Appendix 3 Figure 1.**
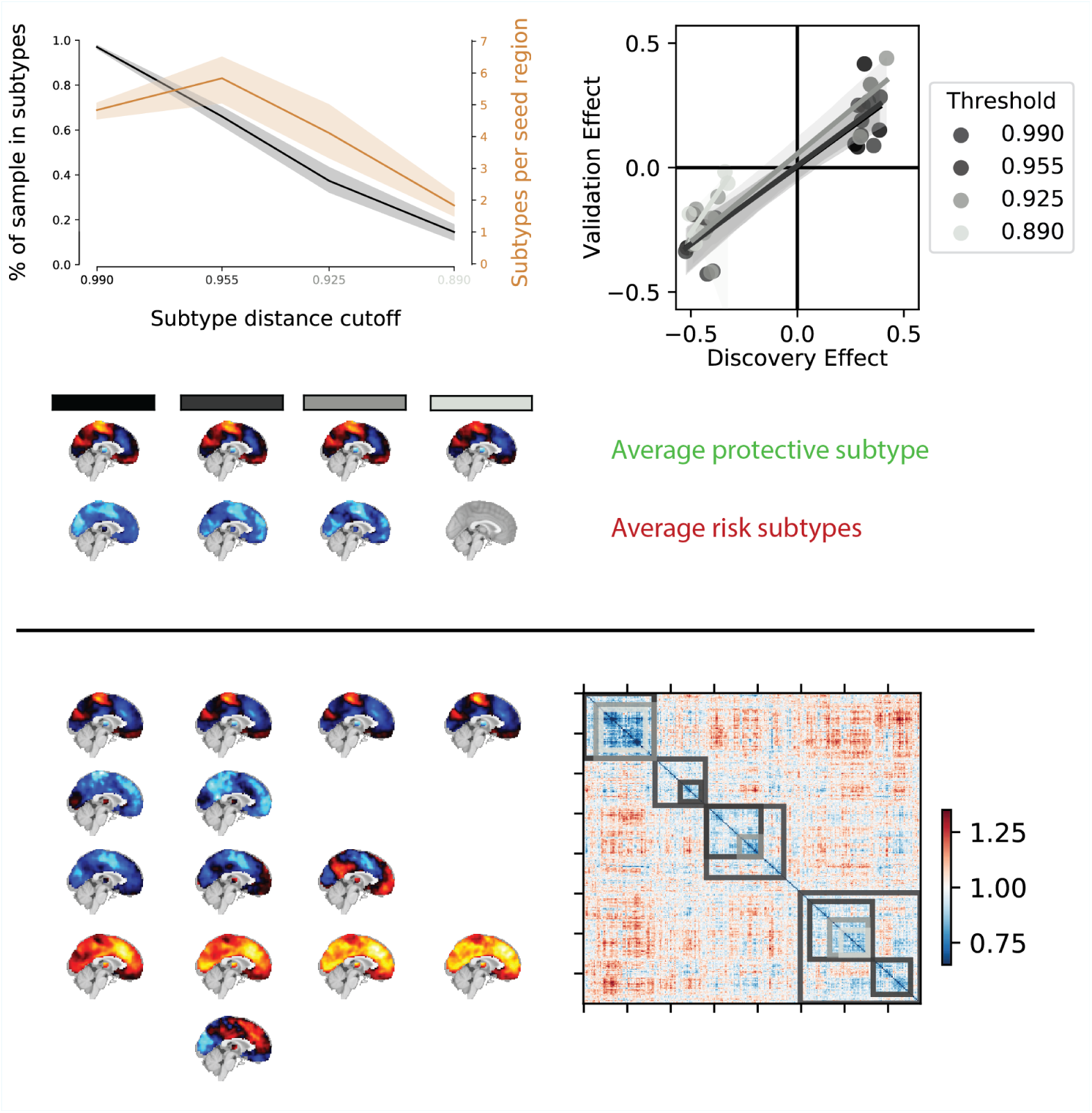
Overview of the robustness of subtype associations with ASD diagnosis to changes in the distance cutoff parameter. a) Average percentage of sample assigned to any subtype (black line) and average number of identified subtypes (orange line) for different levels of distance cutoff parameters. Shaded areas show range of values across all seed networks. Note that both the number of subtypes and the average percentage of the sample described by these subtypes sharply drops off for increasingly stringent FC distance thresholds (black to light grey shaded values on the horizontal axis). b) Correlation of the effect size of subtype association with ASD diagnosis in the discovery and replication data for different levels of distance threshold parameters. Lighter colors reflect more stringent distance cutoff thresholds. Note that the reproducibility of the observed effect sizes remains largely unaffected for small changes of the distance threshold parameters. c) Average subtype maps of protective (top row) and risk (bottom row) for different levels of cutoff parameters. The shaded squares correspond to distance cutoff levels in a) and b). Note that the spatial pattern of the average subtype maps are highly preserved across different thresholds. d) Breakdown of subtype maps across different levels of thresholds illustrated by the example of the dorsal somato-motor seed network. Rows correspond to subtypes and columns correspond to threshold levels. Note that the spatial pattern of individual subtype maps are highly preserved across increasingly stringent threshold levels and that subtypes are rarely split into subsets but rather the total number of subtypes is reduced (x denotes subtypes removed by an increase in the distance threshold). e) Breakdown of subtypes across threshold levels illustrated by the example of the subject by subject dissimilarity matrix of the dorsal somato-motor seed network. Grey shaded overlays reflect the subtype solutions at different dissimiliarity thresholds.

